# Estimation of animal location from grid cell population activity using persistent cohomology

**DOI:** 10.1101/2023.01.10.523361

**Authors:** Daisuke Kawahara, Shigeyoshi Fujisawa

## Abstract

Many cognitive functions are represented as cell assemblies. For example, the population activity of place cells in the hippocampus and grid cells in the entorhinal cortex represent self-location in the environment. The brain cannot directly observe self-location information in the environment. Instead, it relies on sensory information and memory to estimate self-location. Therefore, estimating low-dimensional dynamics, such as the movement trajectory of an animal exploring its environment, from only the high-dimensional neural activity is important in deciphering the information represented in the brain. Most previous studies have estimated the low-dimensional dynamics behind neural activity by unsupervised learning with dimensionality reduction using artificial neural networks or Gaussian processes. This paper shows theoretically and experimentally that these previous research approaches fail to estimate well when the nonlinearity between high-dimensional neural activity and low-dimensional dynamics becomes strong. We estimate the animal’s position in 2-D and 3-D space from the activity of grid cells using an unsupervised method based on persistent cohomology. The method using persistent cohomology estimates low-dimensional dynamics from the phases of manifolds created by neural activity. Much cognitive information, including self-location information, is expressed in the phases of the manifolds created by neural activity. The persistent cohomology may be useful for estimating these cognitive functions from neural population activity in an unsupervised manner.

**Author summary:** Hippocampal place cells fire only when the animal is in a specific position in the environment. Grid cells in entorhinal cortex fire to spatial locations in a repeating hexagonal grid. Information about self-location in the environment is expressed by the population activity of place cells and grid cells. The brain cannot directly observe the information of self-position in the environment but relies on the direction of movement, distance, and landmarks to estimate self-position. This corresponds to unsupervised learning. Estimating the position of an animal from neural activity alone, without using information about the animal’s position, is important for understanding the brain’s representation of information. Unsupervised learning methods using artificial neural networks and Gaussian processes have been used in previous studies to address this problem. However, we show that these previous studies cannot estimate the position of an animal in two dimensions from the population activity of grid cells. As an alternative to the previous studies, we used a topological method called persistent cohomolohy to estimate the animal’s position in 2D and 3D space from the population activity of grid cells. However, it was impossible to estimate the animal’s position from the population activity of place cells. We discussed the causes and solutions to this problem.

## Introduction

In neuroscience, advances in measurement techniques have made it possible to simultaneously record more than 700 neurons in silicon probes [1] [2] and more than 16,000 neurons in two-photon microscope [3]. Behind the high-dimensional neural activity, low-dimensional dynamics are hidden [4] [5]. Low-dimensional dynamics contain important information that is not directly observable from higher-dimensional neural activity, such as task variables, the self-position in the environment, and sensory stimulus variables, such as the angle of visual stimuli. Therefore, estimating low-dimensional dynamics from only high-dimensional neural population activity is an important issue in neuroscience.

When an animal moves on a two-dimensional plane, grid cells in the entorhinal cortex fire at multiple positions on a triangular lattice in the two-dimensional plane [6]. Thus, information about self-position in the environment is represented by the population activity of grid cells. In this study, we estimate the animal position in the environment from the population activity of grid cells without using actual animal position information.

In previous studies, unsupervised learning using Gaussian processes or artificial neural networks has estimated low-dimensional dynamics **s** from high-dimensional neural activity **o**. For example, a method using Gaussian processes estimates the rat’s position in its environment, a low-dimensional dynamic, from about 30 place cells recorded from the rat’s hippocampus [7]. Artificial neural network-based methods estimate the angle of visual stimuli, a low-dimensional dynamics, from the activity of 63 neurons recorded from the macaque primary visual cortex, the rat’s position on a one-dimensional linear line from the activity of 100 grid cells generated by the simulations [8], and the monkey’s hand trajectory from the activity of about 200 neurons recorded from the monkey’s motor cortex [4]. The high-dimensional neural activity **o** can be written as **o** = *f* (**s**) using the function *f*. The previous studies mentioned above are for cases where the function *f* is relatively simple. The stronger the nonlinearity of the function *f*, the more difficult it is expected to estimate the low-dimensional dynamics **s** from the high-dimensional neural activity **o**. For example, it is expected to be more difficult to estimate the animal position on a two-dimensional plane rather than on a one-dimensional line from the population activity of grid cells. Note that the form of the function *f* is unknown here. In this study, we show experimentally and theoretically that previous methods using Gaussian processes and artificial neural networks fail to estimate low-dimensional dynamics as the nonlinearity of the function *f* becomes strong.

A method that applies topology called persistent cohomology [9] is another approach to Gaussian processes and artificial neural networks. We describe an overview of persistent cohomology below. In persistent cohomology, the topological structure of the data is examined in an n-dimensional space in which high-dimensional (n-dimensional) neural activity data are distributed. We consider each data point as an n-dimensional sphere and gradually increase the sphere’s radius (Fig1A). The graph in Fig1B is called barcode. The horizontal axis of the graph represents the radius of the n-dimensional sphere. The vertical axes H^0^, H^1^, H^2^ denote 0-dimensional, 1-dimensional, and 2-dimensional holes, respectively. The length of the green line in the barcode indicates how long the hole in the data structure persists as the radius of the n-dimensional sphere is increased. As the radius of the n-dimensional sphere of data points is gradually increased, all the data points overlap to form a one-dimensional hole H^1^, indicated by the red circle (Fig1A). The radius of the n-dimensional sphere at this point is the start of the green line indicated by the arrow of H^1^ (Fig1B). As the radius of the n-dimensional sphere is further increased in Fig1A, there is a timing when the 1-dimensional hole H^1^ disappears, as shown by the last arrow, and at this time, the green line indicated by the arrow of H^1^ is broken (Fig1B). The longer the barcode line, the more the hole reflects the topological features of the data structure. Short barcode lines are a weak topological feature of the data structure and reflect the noise in the data. Depending on how many holes H^0^, H^1^, H^2^ in each dimension have long barcode lines, we can find the topological data structure (Fig1C). In Fig1B, the long lines indicated by the arrows are H^0^ = 1, H^1^ = 1, H^2^ = 0, and we can find that the topological structure of the data is ring-shaped (Fig1C).

**Fig 1.**
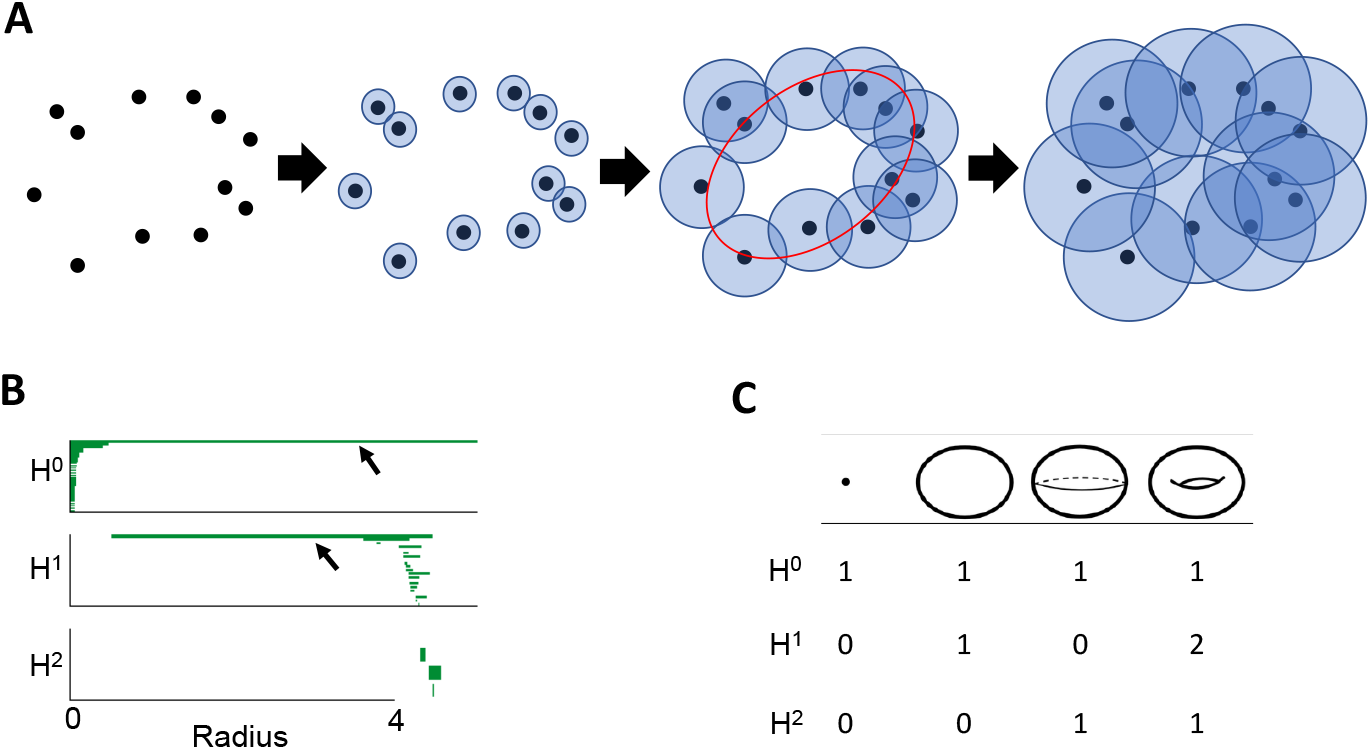
Schematic pictures of persistent cohomology. (A) Conceptual diagram of the change in data structure as the sphere radius of the data points increases. Each data point corresponds to neural activity at each time point. (B) Barcode. H^*n*^ denotes the n-dimensional hole created by the set of data spheres. As the radius of the data spheres is increased, there is a time when the n-dimensional hole appears, which is the start time of the green line. The length of the green line corresponds to the duration of the n-dimensional hole as the radius of the data spheres increases. (C) Example of topological classification of data structures based on the number of n-dimensional holes H^*n*^. The numbers in the table are called betti numbers (Materials and methods).

H^*n*^ contains the phase information of n-dimensional holes. We estimate the low-dimensional dynamic from the phase information of the manifold with n-dimensional holes formed by the high-dimensional neural activity. However, persistent cohomology does not allow us to estimate low-dimensional dynamics from any high-dimensional neural activity. As shown in the Results, persistent cohomology cannot estimate the animal’s location from the population activity of place cells. The method using persistent cohomology is valid only when low-dimensional dynamics are reflected on the phase of the manifold formed by high-dimensional neural population activity.

In previous studies, persistent cohomology has found a ring-shaped data structure of H^0^ = 1, H^1^ = 1, H^2^ = 0 in the population activity of place cells and head orientation cells [10] [11] [12] [13] [14]. In particular, in the case of head direction cells, the animal’s head direction can be estimated from 0 to the 360-degree phase of the ring-shaped data structure. Also, persistent cohomology has revealed a 2-dimensional torus structure H^0^ = 1, H^1^ = 2, H^2^ = 1 in the activity of about 150 grid cells recorded from a rat exploring a two-dimensional plane [5]. It was also found that the positional information of the animal in 2-D space is reflected in the phase of the 2-D torus. Based on this fact, Kang et al. estimated the animal’s position in the 2-D plane by applying persistent cohomology to the activity of grid cells generated by the simulation [15]. We estimate the rat’s position on a 2-D plane from the actual grid cell population activity.

Grid cells in rat and human brains are also found in 3D space [16] [17]. However, it is not known what the topological data structure is for the population activity of grid cells while an animal moves through 3D space. In this study, we also address this issue through simulations. By applying persistent cohomology to the population activity of grid cells generated by simulation, we found a 3D torus as a low-dimensional dynamics. Furthermore, we estimated the animal position in 3D space from the phase information of the three rotational axes of the 3D torus.

## Results

First, we pointed out the problems of previous methods using Gaussian processes and artificial neural networks through simulation experiments, using place cells and grid cells as examples, respectively. Next, we used persistent cohomology to estimate the animal position in 2-D and 3-D space from the population activity of grid cells.

### Estimation of animal location from place cells using previous methods

First, we estimate the animal location from place cell activity by the previous methods, PLDS [18], PfLDS [8], LFADS [4], and P-GPLVM [7]. A detailed explanation of these methods can be found in Materials and methods.

We outline how the previous methods estimate low-dimensional dynamics from high-dimensional neural activity. The estimation is unsupervised learning and consists of an encoder and a decoder. Given high-dimensional neural activity **o**, the encoder outputs an estimate of low-dimensional dynamics as ŝ = *g*_*ϕ*_(**o**). Here, the function *g*_*ϕ*_ is the encoder model of the artificial neural network or Gaussian process, and *ϕ* is the parameter of the encoder model. The decoder outputs **ô** = *g*_*ψ*_(ŝ), which reconstructs the given high-dimensional neural activity from ŝ estimated by the encoder. The function *g*_*ψ*_ is the decoder model of the artificial neural network or Gaussian process, and *ψ* is the parameter of the decoder model. We train the model’s parameters to reduce the error between the high-dimensional neural activity **o** and the decoder outputs **ô**. By using the model’s parameters after the learning has converged, we obtain an estimated low-dimensional dynamics.

Fig 2A left shows the animal’s trajectory on a 2D plane for 5000 s. Fig 2A right shows the receptive field of a place cell. Fig 2B shows the activity of 100 place cells for 5000 s. Bin size is 1 s for all the following experiments. Fig 2C shows the estimation of animal location by the previous methods.

**Fig 2.**
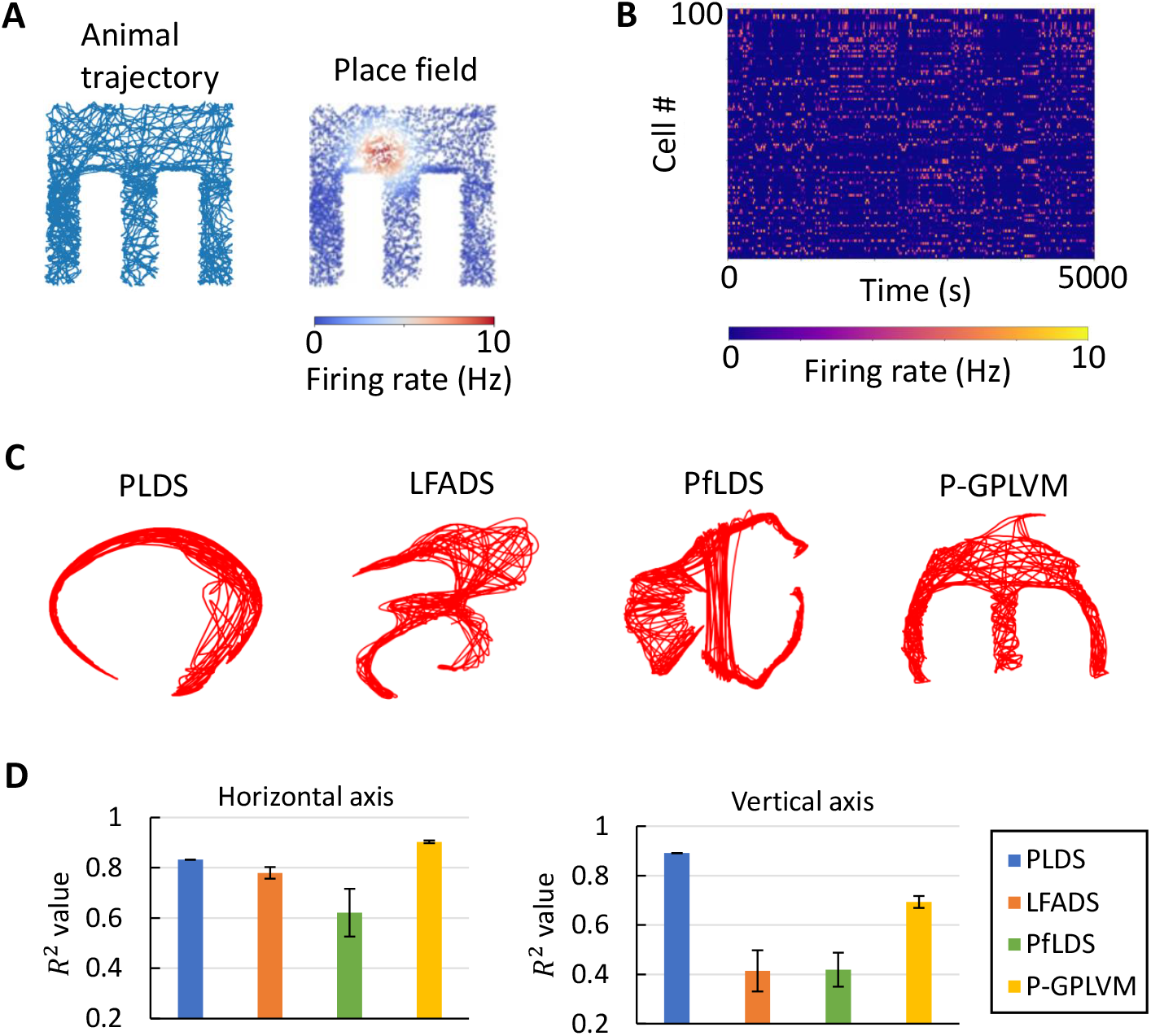
Estimation of animal location from place cells by artificial neural networks and Gaussian processes. (A left) Animal trajectory for 5000 seconds. (A right) A receptive field of a place cell. Other receptive fields of 100 place cells are shown in Fig **??**. (B) Raster plots of 100 place cells for 5000 seconds (C) Results of animal location estimation using previous methods. Here we show the best estimation results out of 6 experiments. We changed the initial values of the model’s parameters in each experiment. See Materials and methods for details. (D) *R*^2^values with error bars in 6 experiments.

Fig 2D shows the *R*^2^ value calculated using the estimation results of the four previous methods and the actual animal location. The *R*^2^ value is defined as follows

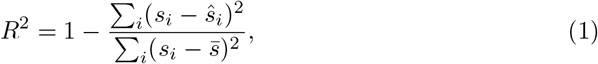

where *s*_*i*_ is the position of the animal at time *t*_*i*_, ŝ_*i*_ is the estimated position of the animal at time *t*_*i*_, and 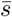 is the average of the animal’s positions. The closer *R*^2^ is to 1, the closer the estimated value is to the actual animal location.

### Estimation of animal location from grid cells using previous methods

P-GPLVM could estimate animal location to some extent for place cells (Fig 2C). Next, using the previous method, we estimate the animal’s location from the grid cells’ activity. Grid cells have more complex receptive fields than place cells (Fig 3A), which would make it more difficult to estimate the animal’s location from neural activity.

**Fig 3.**
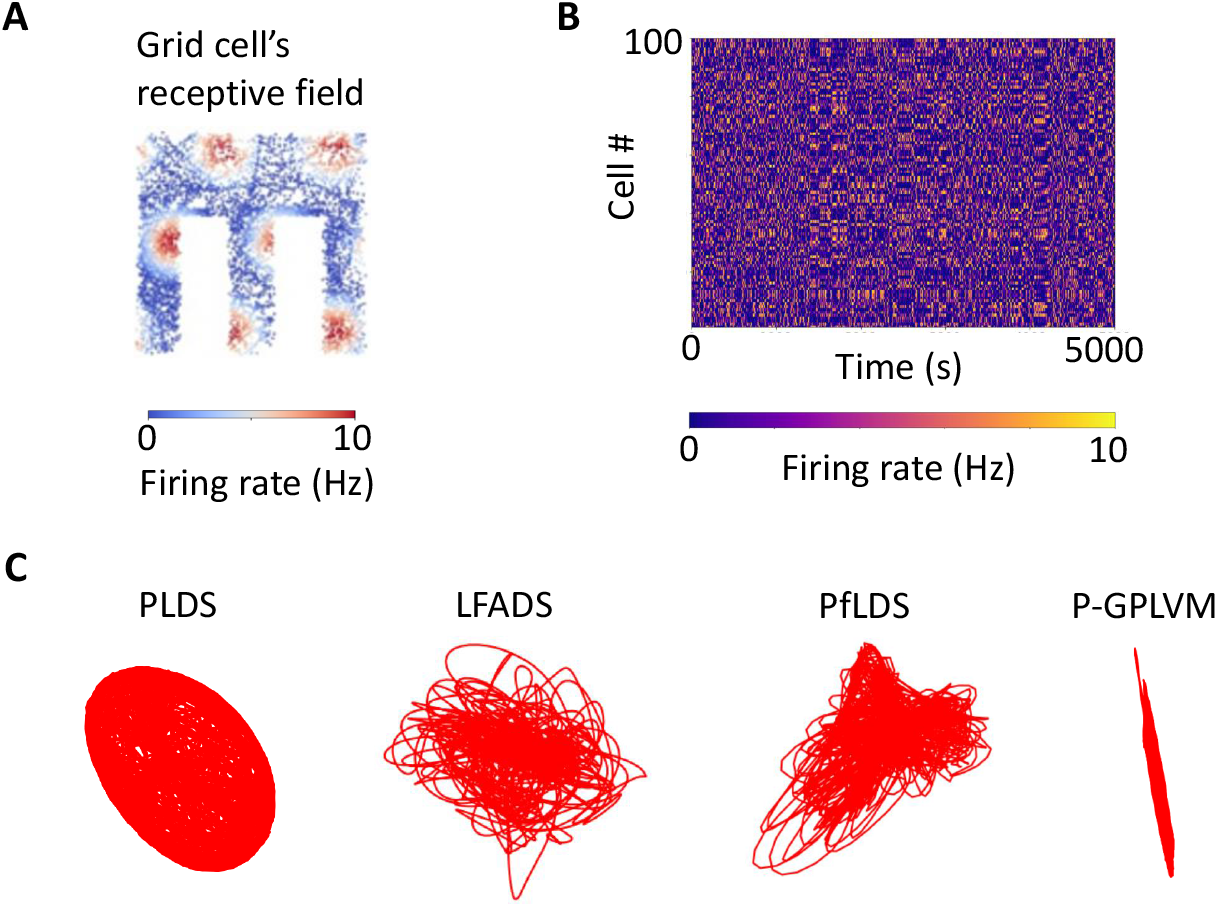
Estimation of animal location from grid cells by artificial neural networks and Gaussian processes. (A) Receptive field of a grid cell. Other receptive fields of 100 grid cells are shown in S2 Fig. (B) Raster plots of 100 grid cells for 5000 seconds. (C) Estimation of animal location by previous methods. Here we show the best estimation results out of 6 experiments. We changed the initial values of the model’s parameters in each experiment. The hyperparameters of each model are the same as in the place cell experiment. See Materials and methods for details.

Fig 3B shows the activity of 100 grid cells for 5000 seconds. Fig 3C shows the estimation of animal location by the previous methods. All four previous methods are not successful in estimating the animal location compared to place cells. This may be because the receptive fields of grid cells are more complex than those of place cells, making it more difficult to model the function *f* of **o** = *f* (**s**). Another problem of previous methods is that even if the error between the model’s decoder output **ô** and the given high-dimensional neural activity **o** is close to zero, the estimated low-dimensional dynamics ŝ is not guaranteed to be close to the true low-dimensional dynamics **s**. This is because even if we estimate low-dimensional dynamics that are completely different from the true low-dimensional dynamics, the decoder function *g*_*ψ*_ can output **ô** that has an error close to zero with the given high-dimensional neural activity **o**.

### Estimation of animal location from grid cells using persistent cohomology

Previous methods using artificial neural networks and Gaussian processes do not work well for estimating animal positions from grid cells. In the following, we use persistent cohomology to estimate the position of animals in 2-D and 3-D space from grid cells.

### Estimation of animal location in 2D space

We show the results of applying persistent cohomology to 100 grid cells in Fig 3A.

As shown by the arrows in Fig 4A, the long lines of the barcode indicating the features of the data structure are H^0^ = 1, H^1^ = 2, H^2^ = 1, that is, a two-dimensional torus structure (Fig 1C), which can be visualized in three-dimensional space using two parameters *t, p* (0 ≤ *t* ≤ 2*π*, 0 ≤ *q* ≤ 2*π*) by the following.

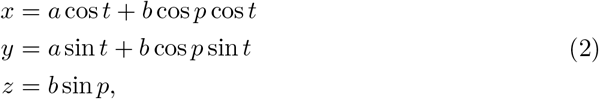

**Fig 4.**
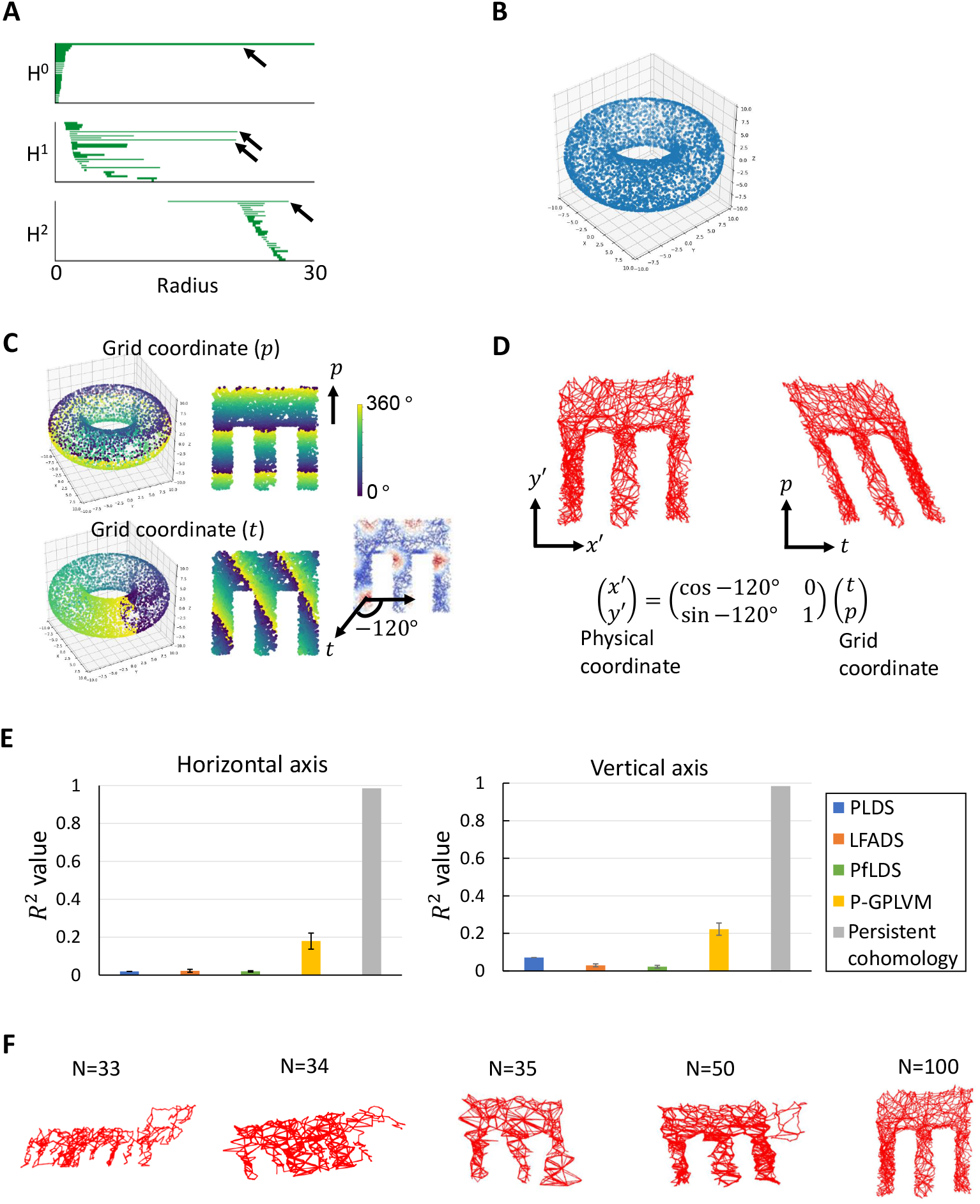
Estimation of animal location from grid cells using persistent cohomology. (A) Barcode. (B) 2D torus structure. Each point corresponds to ℝ^100^→ ℝ^3^ neural activity at each time. (C) Torus phase 0 ∼360 degrees and animal position at each time. (D left) Estimated position after transformation to Cartesian coordinate, or physical coordinate. (D right) Animal position estimated from the torus phase information. The axis *t, p* of the torus created by the activity of grid cells is not Cartesian in the physical world but oblique coordinates, as shown in (C). Therefore, if *t, p* is set to Cartesian coordinates, the estimation results become oblique. (E) *R*^2^ values with error bars. For the method using persistent cohomology, we show the result for N=100 in sparse circular coordinates. (F) The result of sparse circular coordinates. N is the number of subsets of 5000 data points used to obtain low-dimensional coordination.

where *a, b* are the radius of the large and small circles of the 2D torus *S*^1^ ×*S*^1^, respectively. See Materials and methods for how to obtain the parameters *t, p* from persistent cohomology. Fig 4B shows a 2-D torus, which is the low-dimensional dynamics obtained from a high-dimensional (n=100) grid cell activity. Each point in the torus corresponds to neural activity at each time point per second. The phase of the torus corresponds to animal location and the scale of the grid pattern of the grid cell’s receptive field (Fig 4C).

Next, we considered estimating the animal position from the phase of the torus. However, there are two problems with the estimation. The first problem is that the animal position is not uniquely determined due to the periodicity of the torus (Fig 4C). Therefore, to eliminate the periodicity, we added 360 degrees to the current phase of the torus for each round of the torus and subtracted 360 degrees for each round of the torus in the opposite direction. The second problem is that the coordinate system created by the parameters *t, p* is not a Cartesian coordinate system. The direction in which the parameter *t* moves is tilted − 120 degrees from the horizontal axis (Fig 4C). Therefore, if the parameters *t, p* is considered in the Cartesian coordinate system, the estimated animal movement trajectory will be tilted (Fig 4D right). Therefore, we used a matrix to transform the coordinate system created by the parameters *t, p* into a Cartesian coordinate system (Fig 4D bottom). Through these operations, we could accurately estimate the animal’s location (Fig 4D left). The previous methods using artificial neural networks and Gaussian processes could not estimate the animal position on a two-dimensional plane from grid cells. In contrast, the persistent cohomology method could accurately estimate the animal position (Fig 4E). In addition, by using a technique called sparse circular coordinates [38](See Materials and methods), persistent cohomology method can obtain low-dimensional coordinates from only a subset of data points (Fig 4F).

Next, we examined the correspondence between the phase of the torus and the scale of the grid pattern of the receptive field of grid cells. The grid pattern’s scale in the grid cell’s receptive field decreases from ventral to dorsal in the entorhinal cortex [6]. We applied persistent cohomology to the neural activity of 100 grid cells for each small and large grid pattern scale. We confirmed that the phase of the torus corresponds the scale of the grid pattern of the receptive field (Fig 5). The directions of phase parameters *p, t* of the torus correspond to the orientation of the traveling wave of the grid pattern.

**Fig 5.**
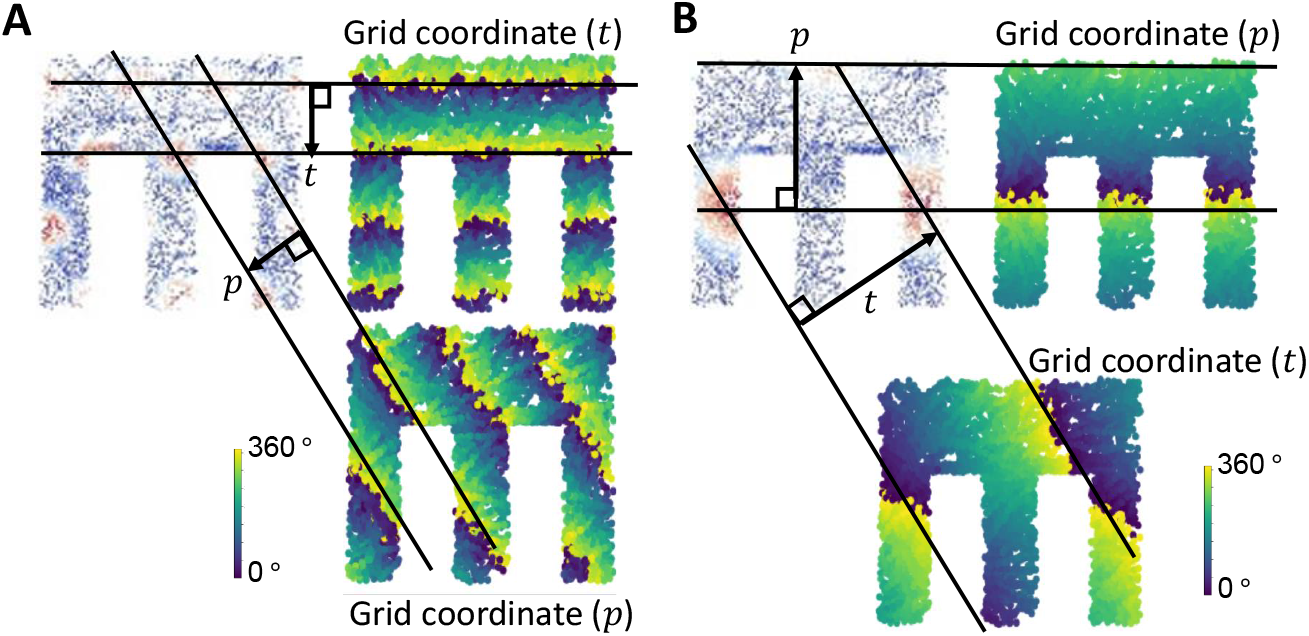
Correspondence between the parameters *p, t* of a 2-D torus and the grid pattern. Regardless of the grid pattern scale, the axis *t, p* of the 2D torus created by the grid cells’ activity corresponds to the direction of the wavefront of the grid pattern indicated by the parallel lines. Other receptive field of grid cells is shown in S3 Fig. (A) Small grid scale. (B) Large grid scale.

Finally, we estimated the rat’s position from the neural activity of 149 grid cells recorded from the rat’s entorhinal cortex during exploration on the 2-D plane [5](S5 Fig). We confirmed by persistent cohomology that the population activity of grid cells forms a 2-D torus and that the torus phase reflects the rat’s positional information (Fig 6A). We estimated the rat’s position from the torus phase using the same procedure as in the simulation (Fig 6B). In y-axis, the estimated result is smaller than the actual rat movement width, but the timing of the vertical movement is almost identical. In the x-axis, the estimated results almost coincide with the actual movement trajectory of the rat. One possible reason for the discrepancy in y-axis is that the actual grid pattern of the rat’s gird cell is a little distorted than the grid pattern of grid cells generated by the simulation (S5 Fig **??**). Like the simulations, persistet cohomology method was able to estimate the rat’s location more accurately than previous methods(Fig 6C).

**Fig 6.**
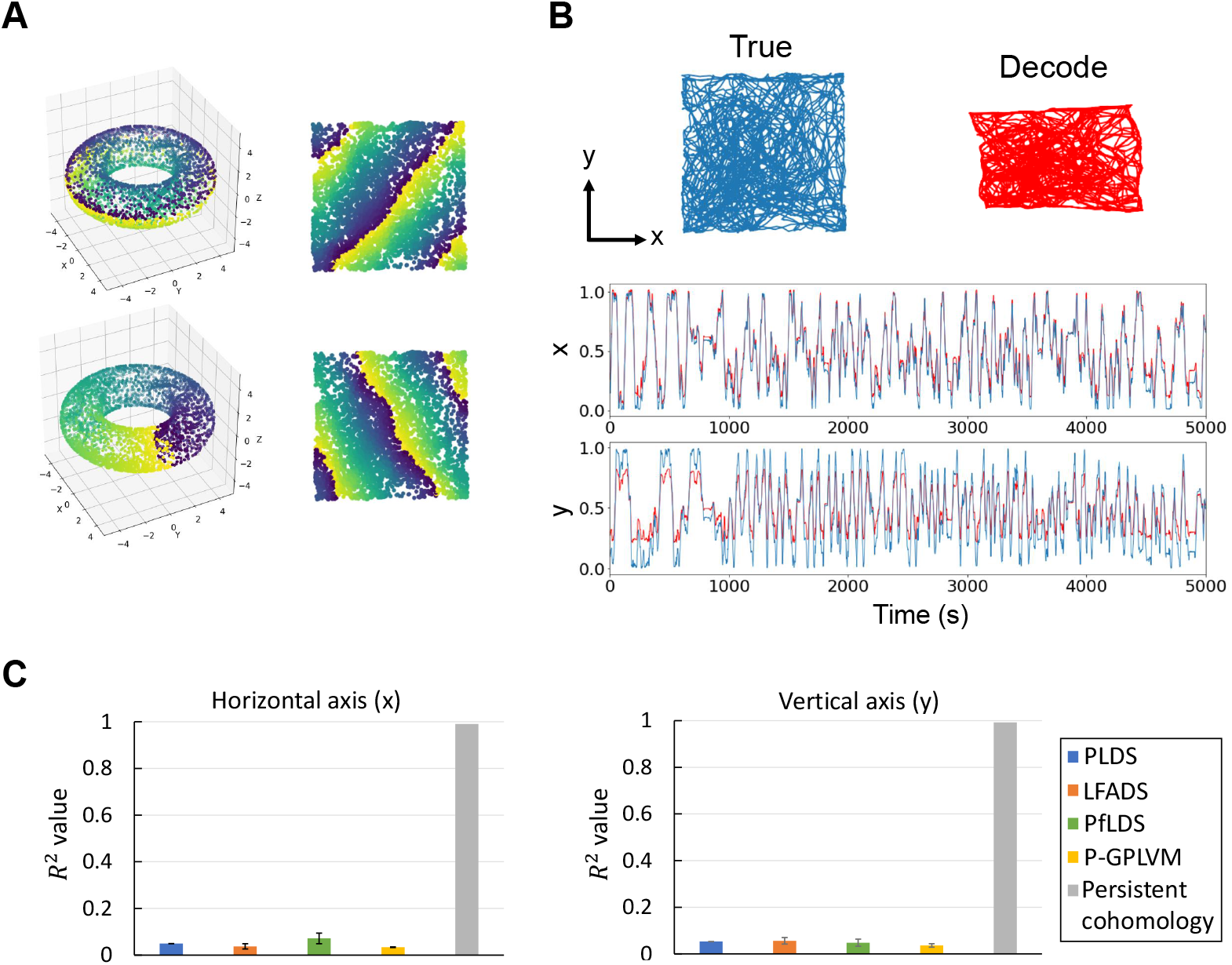
Estimation of rat’s location from the neural activity of 149 grid cells recorded from the rat’s entorhinal cortex using persistent cohomology. (A) Torus phase and animal location at each time point. (B) Comparison of actual rat’s trajectory (blue) and estimated rat’s trajectory (red). (C) *R*^2^ values with error bars. The experimental conditions are the same as in the earlier experiment except for the input data.

### Estimation of animal location in 3D space

Experiments using fMRI have reported that the activity of grid cells in the entorhinal cortex when humans are exploring in 3D space is a face-centered cubic lattice [17].

Therefore, we generated by simulation the activity in 20000 s of 3000 grid cells firing at the positions of a face-centered cubic lattice. Fig 7A shows the raster plots of gird cells’ activity. Fig 7C shows the receptive field of one grid cell. After applying persistent cohomology to the high-dimensional neural activity, we found H^0^ = 1, H^1^ = 3, H^2^ = 3, H^3^ = 1 (Fig 7B). This is topologically equivalent to a 3-dimensional torus [19] [20].

**Fig 7.**
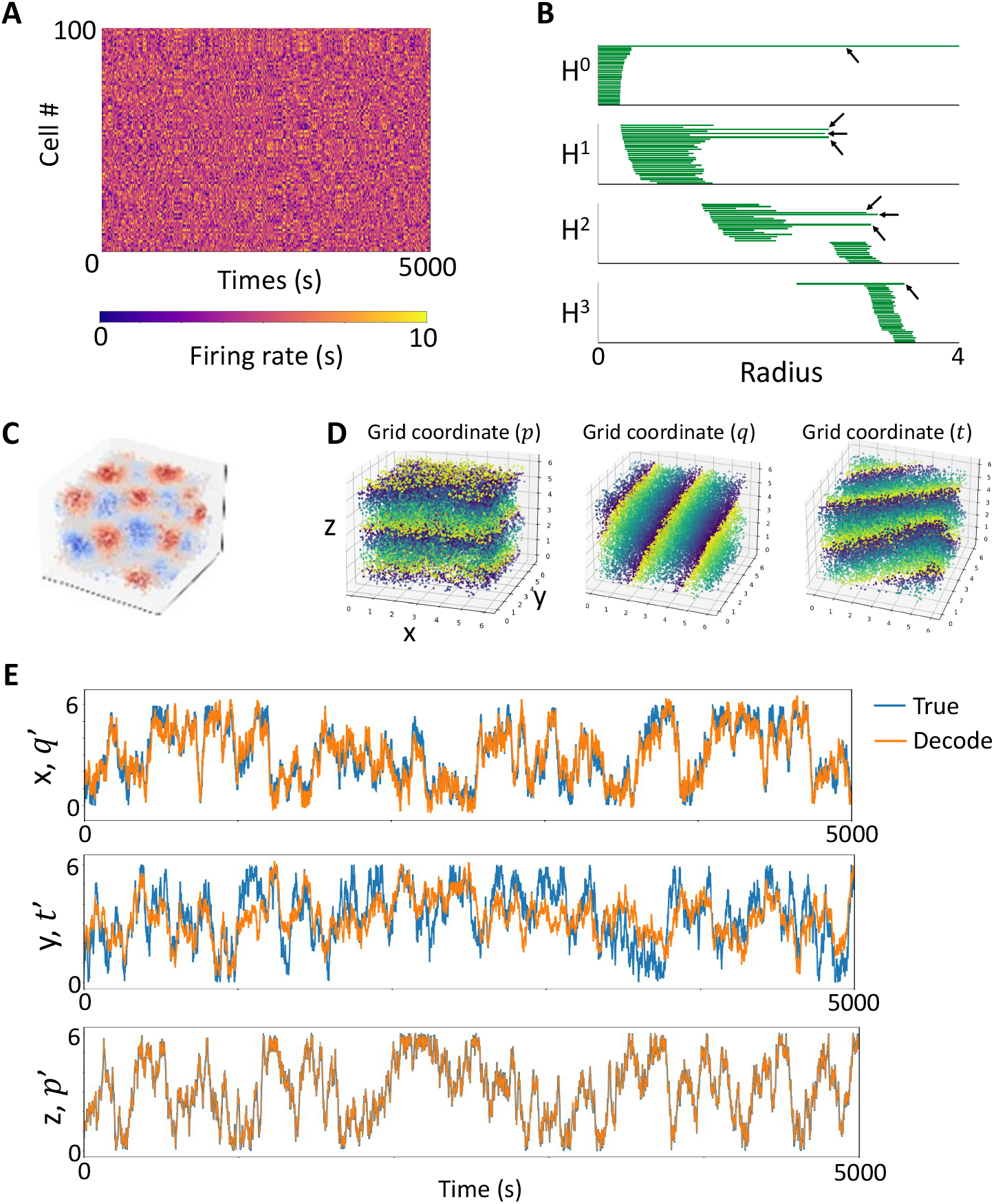
Estimation of animal location in 3D space using persistent cohomology. (A) Raster plots of 100 grid cells for 5000 seconds. (B) Barcode. H^0^ = 1, H^1^ = 3, H^2^ = 3, H^3^ = 1. (C) A receptive field of a grid cell in 3D space. The receptive fields of other grid cells are shown in S6 Fig. (D) Correspondence between the phase of 3D torus and the animal’s location. S1 video for more detail. (E) Comparison of actual animal location (blue) and estimated location (orange). N=1000 in sparse circular coordinates.

The phase of the 3D torus reflects the animal positions in 3D-space (Fig 7D). The coordinate system *p* is oriented -180 degrees to the *z* axis, the coordinate system *q* is oriented 150 degrees to the *x* axis, and the coordinate system *t* is oriented -150 degrees to the *y* axis (Fig 8). Therefore, we used the following coordinate transformations to align the angles of the coordinate systems.

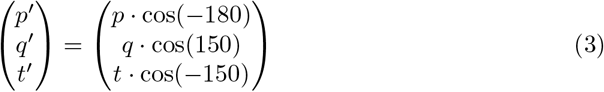

**Fig 8.**
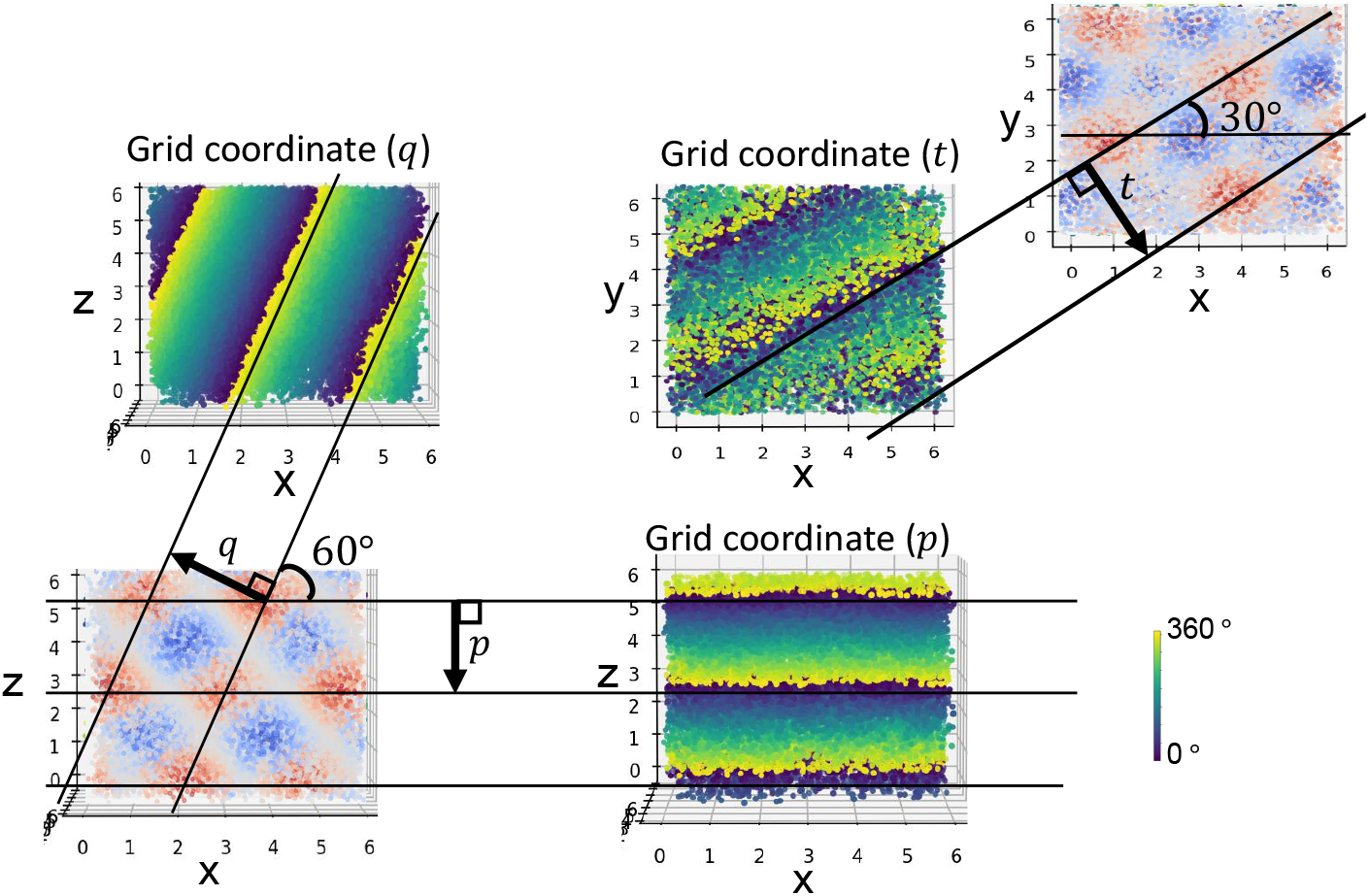
Correspondence between the coordinate system of the paratemers *p, q, t* of the 3-D torus and the grid pattern. Like in 2-D, the axis *p, q, t* of the 3-D torus created by the grid cells’ activity corresponds to the direction of the wavefront of the grid pattern indicated by the parallel lines.

As in the 2D case, for each of the three parameters *t, p, q* of the 3D torus, we performed a 360-degree addition operation for each round of the torus and a 360-degree subtraction operation for each round in the opposite direction. Moreover, we obtained the parameters of scaling *a* and translation *b* to align the *t, p, q* with the animal trajectory coordinate **x, y, z** by minimizing the following equation.

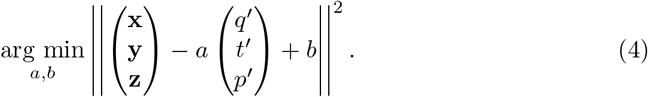

Fig 7E shows the animal position obtained from the phases of the 3D torus. The estimated positions of the x-axis and z-axis are almost identical to the actual positions of the animals, while the y-axis deviation was larger because the parameter *t* and the y-axis position of the animals did not correspond well (Fig 8).

### Applying persistent cohomology to the population activity of place cells

We applied persistent cohomology to the population activity of place cells. We demonstrate by experiment that, unlike in grid cells case, it is difficult to estimate an animal’s location from place cells using persistent cohomology. Fig 9 (A) shows the movement trajectory of the animal for 10,000 seconds within four different forms of environment. Fig 9 (C) shows the results of dimensional reduction of 100 place cell activities into 3-dimensional space by UMAP [21]. The manifold formed by place cells activity reflects the shape of the environment. From the barcode, we see that H^0^ = 1, H^1^ = 0, H^2^ = 0 for the square and E-shape arenas, H^0^ = 1, H^1^ = 1, H^2^ = 0 for the O-shape arena, and H^0^ = 1, H^1^ = 2, H^2^ = 0 for the 8-shape arena (Fig 9 (B)). In the 8-shape arena, there are large and small holes, and this is to examine whether the hole size is reflected in the manifold created by the population activity of place cells. In this experiment, there was little difference in the H^1^ duration of the barcode reflecting the large and small holes. Therefore, we did not find detailed information about the shape of the environment in the manifold created by the population activity of place cells. In summary, the manifolds formed by place cell population activity do not reflect self-location information, but rather reflect the topological shape of the environment.

**Fig 9.**
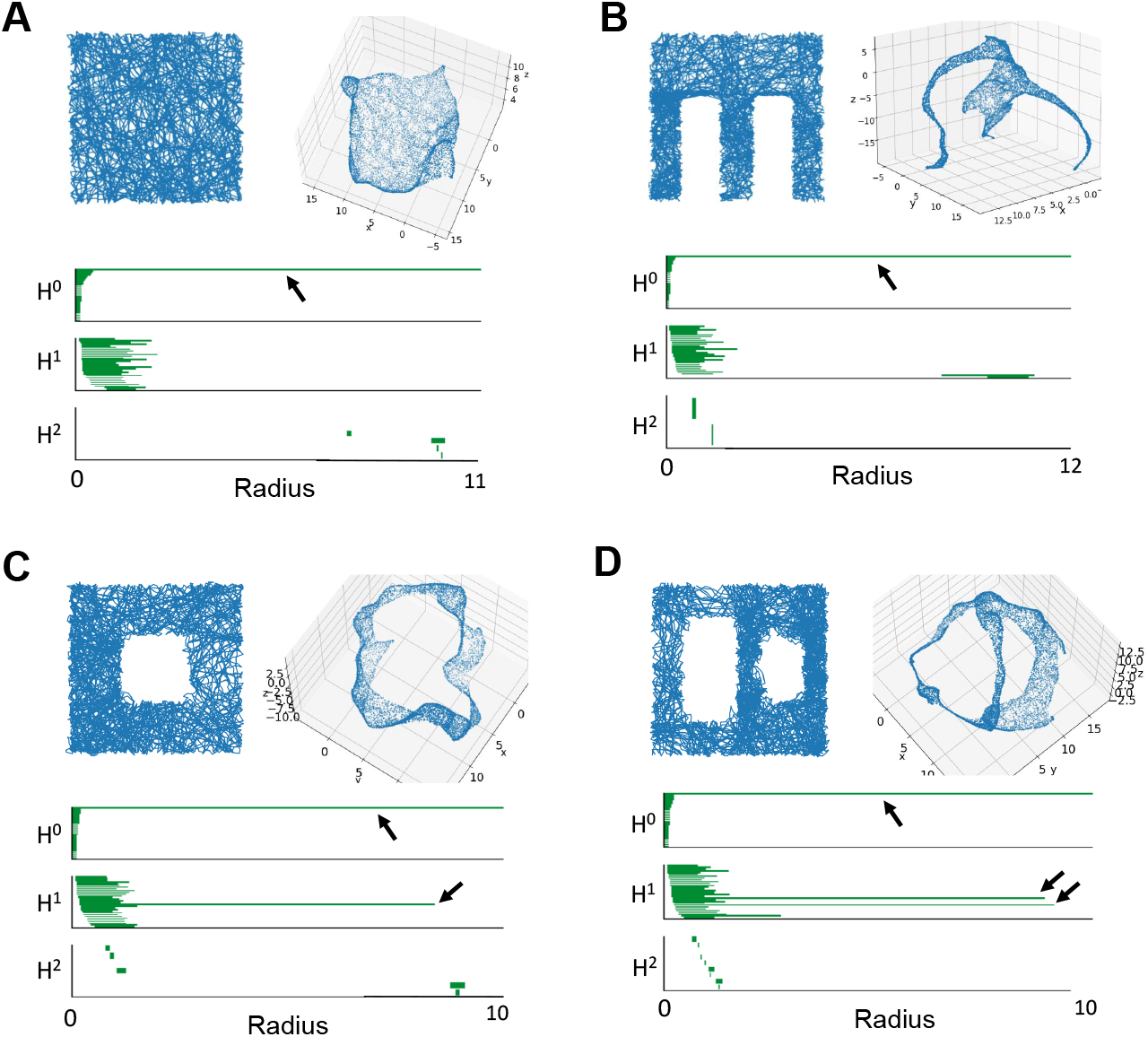
Applying persistent cohomology to the population activity of place cells. We can see that the topological shape of arenas is reflected in the manifold created by place cells’ activity. (A) Square-shape arena. (B) E-shape arena. (C) O-shape arena. (D) 8-shape arena.

## Discussion

The main contributions of this study are listed below.

- We showed experimentally and theoretically that the stronger the nonlinearities between high-dimensional neural activity and low-dimensional dynamics, the less successful the estimation of animal location becomes with conventional methods based on Gaussian processes and artificial neural networks.
- Using persistent cohomology, we estimate the rat’s position in a two-dimensional plane from the actual grid cells’ population activity.
- In the simulation, we found by persistent cohomology that the population activity of grid cells of an animal moving in 3D space results in a 3D torus structure, and we estimated the animal’s position in 3D space from the phase information of the 3D torus.
- We clarified the problems in estimating the animal’s position using persistent cohomology from the population activity of place cells through simulation experiments.

### Advantages of persistent cohomology

The persistent cohomology method estimates the low-dimensional dynamics **s** from the phase information of the manifold created by the neural activity data **o**. Our experiments have shown that the previous methods using artificial neural networks and Gaussian processes cannot estimate **s** from **o** when the function *f* (**o** = *f* (**s**)) is strongly nonlinear, as in grid cells. On the other hand, if the information of **s** is embedded in the phase of the manifold created by the neural activity **o, s** can be estimated using persistent cohomology even if the nonlinearity of *f* is strong.

In this experiment, the dimension of **s** was known in advance as the animal’s location. However, if the dimension of **s** is not known, the method using artificial neural networks or Gaussian processes cannot directly determine the dimension of **s**. On the other hand, the persistent cohomolohy can determine the dimension of **s** from the barcode of the hole dimension H^*n*^ generated by the neural activity.

The computation time to estimate animal positions in persist cohomology is very fast compared to methods using artificial neural networks or Gaussian processes. We used the method of sparse circular coordinates to obtain low-dimensional coordinates from a subset of data points. In Fig 4(E), when the subset of data points was 100 out of 5000 data points, the computation time to obtain low-dimensional coordinates was only 0.24568 seconds. Methods using artificial neural networks or Gaussian processes take about an hour of computation time, and the estimation accuracy is not good. When low-dimensional dynamics are encoded in the phase information of manifolds created by population activities such as grid cells, methods using persistent cohomology outperform previous methods in both computation time and estimation accuracy.

### The problems of persistent cohomology

Although we were able to estimate an animal’s location from the activity of grid cells using the persistent cohomology method, there are still some problems.

The first issue is that, as shown in Results, we could not estimate the location of the animal from place cell activity using the persistent cohomology. The previous method using Gaussian processes have been able to estimate the animal location from place cells’ activity to some extent (Fig 2). It may be possible to determine the detailed shape of the environment by considering the dynamics of the short barcode lines as well as the long barcode lines, which is a subject for future work.

The second issue is that we used the actual animal position information to convert the coordinate system *t, p* of the phase of the 2-D torus into a Cartesian coordinate system. There are four possible angles between the parameters *t, p*: ±120 degrees and ±60 degrees. The transformation matrix to the Cartesian coordinate system was obtained in this experiment based on the animal location information. A possible approach that does not use animal location information is to identify the direction of animal movement by combining it with the activity of the head orientation cells (S7 Fig and S8 Fig).

The last issue is that we could not understand the relationship between the direction of the traveling wave of grid pattern in 3-D space and the direction of the parameters *t, p, q* of the 3-D torus. In the 2-D space, there were four possible angles (±120 and ±60 degrees) between the parameters *t, p*, which corresponded to the direction of traveling waves of the grid pattern. It is a future work to investigate the possible angles between the parameters *t, p, q* in the 3-D space.

### Applications of persistent cohomology

In addition to the representation of animal location, grid cells can also represent social relations [22], the concept of things [23], the visual sense [24] [25] and smell [26], word meaning [27], and mental simulation [28]. This information represented by grid cells is expected to form the dynamics on the torus. Higher-order cognitive functions, such as the concept of objects, may be represented by a torus structure of three or more dimensions. Similar to location estimation in animals, it would be possible to estimate the dynamics of cognition from the phase information of the higher dimensional torus.

In addition, grid cells that represent location information are founded not only in the entorhinal cortex but also in the secondary visual cortex [29], somatosensory cortex [30], cingulate gyrus [31], hippocampal plateau [32], and medial prefrontal cortex [33]. Persistent cohomolohy may help investigate how location information is shared and represented among brain regions.

## Materials and methods

The source code used in the experiment is available in https://drive.google.com/drive/folders/1rDypHfXhtZURXNfZqiWSqahIZaC3Nuw8?usp=sharing. We used ripser for the persistent cohomology library [34].

### Place cells in 2D space

We generated the activity of i-th place cell at animal position *x, y* as

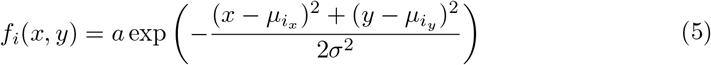

where *a* = 40, *σ*^2^ = 15 in the experiment. We randomly selected *μ*_*i*_, *μ*_*i*_ from a uniform distribution ranging from 0 to *l. l* is the arena’s size, and in this experiment, *l* = 50.

### Grid cells in 2D space

We used continuous attractor neural network model to generate the activity of grid cells by simulation [35]. The activity of a neuron *i* at time *t* + 1, i.e. *f*_*i*_(*t* + 1) is defined as folllowing.

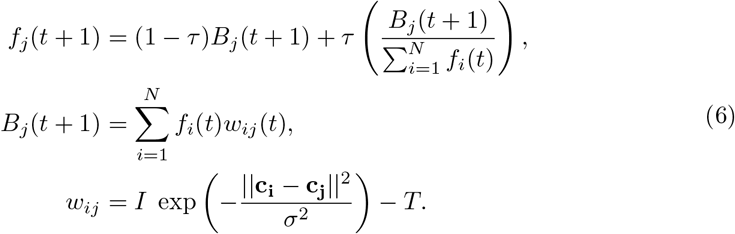

where 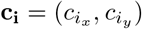 is the position of the cell *i* on 2-D space. *N* is the number of cells in the network. *w*_*ij*_ is the synaptic weight connecting neuron *j* to neuron *i*, with *i, j* ∈{1, 2, …, *N*}. *I* is the intensity parameter, defining the strength of the synapses, regulates the size of the Gaussian and *T* is the shift parameter. The parameter *τ* determines the stabilization strength.

In order to align the coordinate system of the actual rat movement trajectory **x**_*T rue*_, **y**_*T rue*_ with that of the movement trajectory **x**_*Decode*_, **y**_*Decode*_ estimated from the 2-D torus, *a, b, θ*, which adjusts the scaling, translation, and rotation of the coordinate system, was obtained by minimizing the following equation.

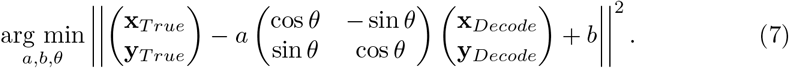

### Grid cells in 3D space

The activity *f* (**r**) of a grid cell firing at a face-centered cubic lattice position **r** in 3-dimensional space was calculated using the following equation [36].

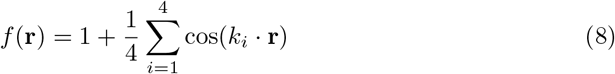

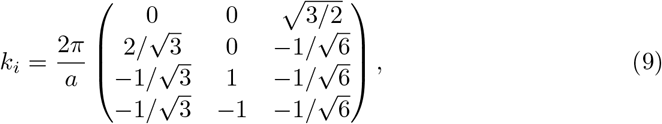

where *a* is the grid size, which was set to *a* = 6 in the experiment. The activity of the 3000 grid cells used in the experiment was generated by substituting **r** + **Δr** into the Eq (8). **Δr** is a random number that follows a uniform distribution between 0∼ 6 on the x,y, and z axes.

### Previous methods

We explain below the previous methods used in our experiments. See the source code (https://drive.google.com/drive/folders/1rDypHfXhtZURXNfZqiWSqahIZaC3Nuw8?usp=sharing) for the hyperparameters of the model used in the experiments.

#### Poisson Linear Dynamical System (PLDS) [18]

PLDS is a generative model based on linear dynamical system (LDS), which uses variational inference to estimate low-dimensional hidden dynamics **s** from neural activity **o**. In LDS, low-dimensional hidden dynamics **s** evolves according to linear Gaussian dynamics:

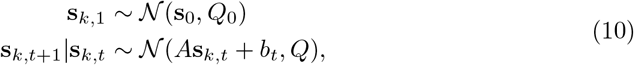

where *k* represents the number of trials in each experiment. In our experiment, we estimated the low-dimensional dynamics from a single trial *k* = 1. Conditioned on **s**, the activity of neuron *i* at time *t* is given by a Poisson distribution

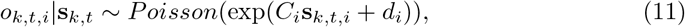

where the matrix C determines how each neuron’s activity is related to the hidden dynamics **s**_*t*_, and the vector *d* represents the mean firing rates of the population activity **o**. PLDS uses an EM algorithm to learn the parameters Θ = {*C, d, A, Q, Q*_0_, *s*_0_ }. The posterior distribution in E-step is given by

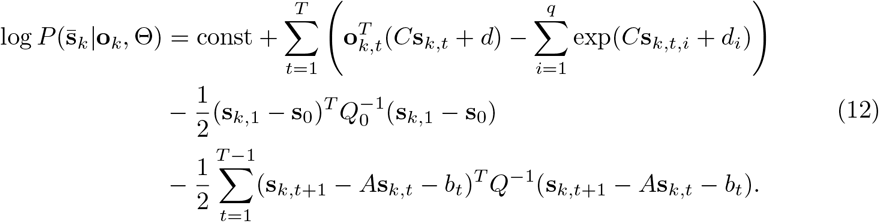

In M-step, we update the parameters Θ by maximizing the following expected joint log-likelihood

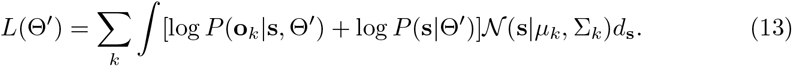

We used a global Laplace approximation [37] to get the mean *μ*_*k*_ and the covariance ∑_*k*_.

#### PfLDS

**[8]:**PfLDS is a modification of PLDS, which models the relationship between low-dimensional dynamics **s** and neural activity **o** using artificial neural networks. In PfLDS, Eq(11) is rewritten as

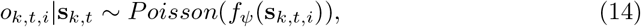

where *f*_*ψ*_ is an arbitrary continuous function from the hidden dynamics **s**_*t*_ into the spike rate. We represent **s**_*t*_ through a feed-forward neural network model. In variational inference, we approximate the intractable posterior distribution *p*_*θ*_(**s** | **o**) by a tractable distribution *q*_*ϕ*_(**s** | **o**) and learn the model parameters *θ* = (*μ*_1_, *Q*_1_, *A, Q, ψ*) by maximizing the evidence lower bound (ELBO) of the marginal log-likelihood:

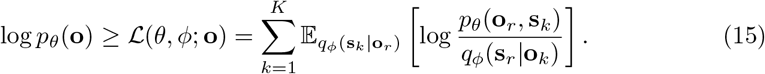

#### Latent Dactor Analysis via Dynamical Systems (LFADS) [4]

In LFADS, the feed-forward neural networks used in PfLDS are replaced by recurrent neural networks (RNNs). We estimate the low-dimensional dynamics **s** by minimizing the following loss function ℒ ^*o*^ − ℒ ^*KL*^

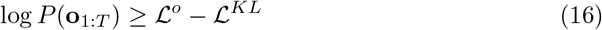

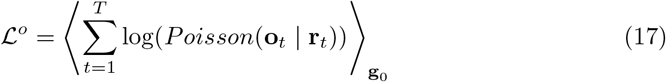

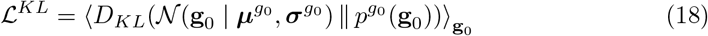

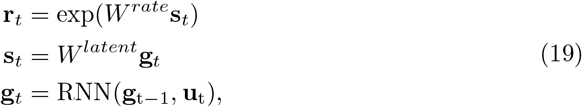

where matrices *W*^*rate*^ maps low-dimensional dynamics **s** to neuron rates **r**, and matrices *W*^*latent*^ maps **g** to **s**. The priors for **g**_0_ and **u**_1_ are diagonal Gaussian distributions. The prior for **u**_*t*_ with *t* > 1 is an auto-regressive Gaussian prior. See [4] for more details.

#### Poisson Gaussian-Process Latent Variable Model (P-GPLVM)

In P-GPLVM, the latent variable dynamics **s** and the tuning curve **f** for the *i* th neuron are modeled by a Gaussian process as follows

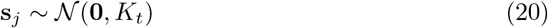

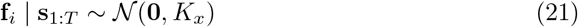

The spike-count of *i*^*′*^ th neuron at *t* given the tuning curve *f*_*i*_ and latent variable dynamics **s**_*t*_ is Poisson distributed as

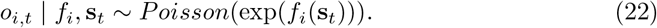

We estimated the low-dimensional dynamics by the following maximum a posteriori (MAP) estimation.

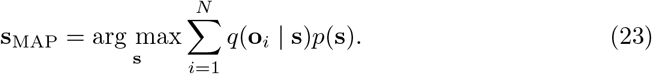

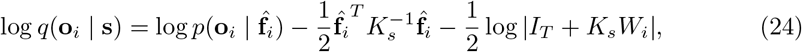

where 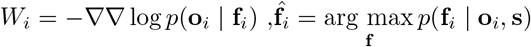, and *I*_*T*_ is the identity matrix (of size *T*).

### Sparse circular coordinates

Sparse circular coordinates is a method for obtaining low-dimensional coordinates using only a subset of the data points *L*⊂ *X* called landmark points in calculating persistent cohomology [38]. We obtained the landmark points by maxmin sampling [39]. First, we randomly select *l*_1_ ∈ *X*. Then, when *l*_1_, *l*_2_, …, *l*_*i−*1_ have chosen inductively, *l*_*i*_ ∈ *X*\{*l*_1_, *l*_2_, …, *l*_*i−*1_} is the data point that maximizes the function

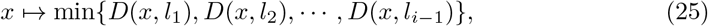

where *D* is the metric space. Continue until the desired number of landmark points are selected. In Fig 4(F), we set the number of landmark points to 33,34,35,50,100 for 5000 data points.

### Homology and cohomology

Persistent cohomology creates barcodes for n-dimensional holes H^*n*^ created by high-dimensional neural activity data. We explain the homology and cohomology required to obtain H^*n*^ [40] [9].

A simplicial complex is a set consisting of *n*-simplex (Fig 10).

**Fig 10.**
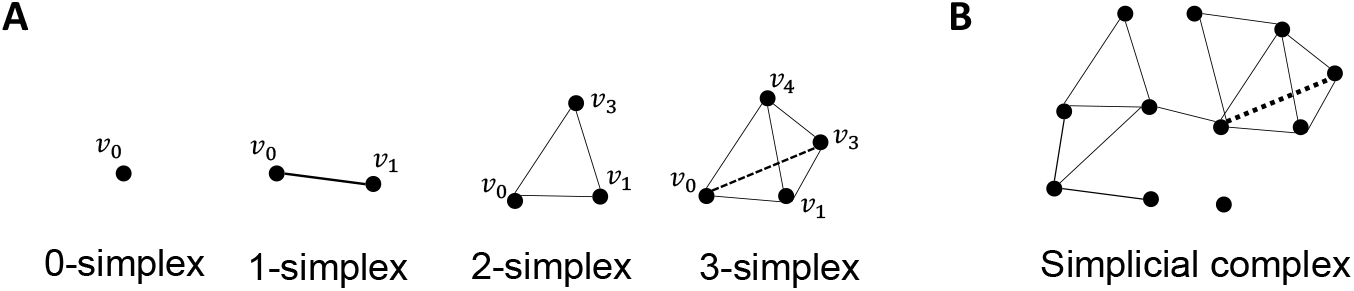
A simplicial complex is a set composed of n-simplex. (A) Example of n-simplex. From right to left: vertices, edges, triangles, and tetrahedron. (B) Example of a single complex.

The boundary of an *n*-simplex *σ* = [*V*_0_, *V*_1_, …, *V*_*n*_] is composed of *n* − 1-simplex combinations. The boundary operator *∂*_*n*_ is defined as follows.

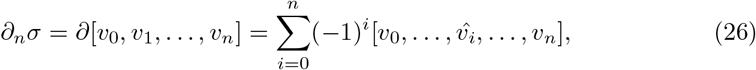

where 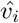 indicates that *v*_*i*_ is deleted from the sequence [*v*_0_, …, *v*_*n*_], that is, *∂*[*v*_0_*v*_1_] = [*v*_1_] − [*v*_0_], *∂*[*v*_0_*v*_1_*v*_2_] = [*v*_1_*v*_2_] − [*v*_0_*v*_2_] + [*v*_0_*v*_1_] for example.

*C*_*n*_, called n-chain groups, is an Abelian group with an *n*-simplex basis and is represented by the following equation with boundary operators.

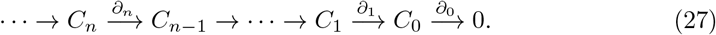

For example, *∂*_1_ : *C*_0_ → *C*_1_ is expressed as

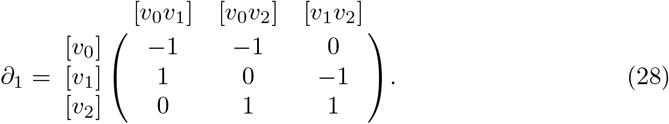

The image (Im) of the boundary operators *B*_*n*_ = Im *∂*_n+1_ is called the boundary group, and kernel (Ker) *Z*_*n*_ = Ker *∂*_n_ is called the cycle group. Then, the homology group H_*n*_ = *Z*_*n*_/*B*_*n*_ is defined.

When *X*^*n*^ is a set of n-simplex and *R* is a commutative ring, the element of *C*^*n*^ = function*f*_*n*_ : {*X*^*n*^ →*R*} is called the co-chain. The coboundary map *d* is expressed as

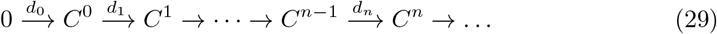

When *Z*^*n*^ = Ker *d*_*n*+1_, the element of *Z*^*n*^ is called the cocycle, and When *B*^*n*^ = Im *d*_*n*+1_, the element of *B*^*n*^ is called the coboundary. Then, the cohomology group H^n^ = Z^n^*/*B^n^ is defined.

### Persistent cohomology

We used the Vietoris-Rips complex (VR) method to construct a single complex from a set of data points of higher dimensional neural activity [41]. Let *P* = *x*_*i*_∈ {ℝ^*N*^ |*i* = 1, …, *l* } be the set of *l* data points in N-dimensional space. In the VR complex, consider an N-dimensional sphere with radius *r* centered at each data point *x*_*i*_ in N-dimensional space. The following equation then defines the VR complex.

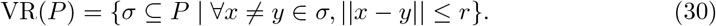

We obtain an increasing sequence of VR complexes VR(*P, r*_1_) → … → VR(*P, r*_*m*_) according to the increasing sequence *r*_1_ *<* … *< r*_*m*_ of radius *r*, which is called filtration. The *r*_*i*_(*i* = 1, …, *m*) is the radius at which the simplex occurred. We obtain the sequence of cohomology for the increasing sequence of VR complexes as follows [42] [9] [40].

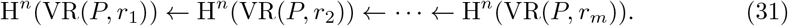

In persistent cohomology, we can create the barcode by examining how long the element of H^*n*^(VR(*P, r*_*i*_)) persists for an increasing sequence of radius *r*. Those with long durations are considered significant topological features of the data structure, while those with short durations are considered noise. The betti number is defined by *b*_*n*_ = dim H^*n*^.

### How to obtain the phase parameters of torus

We describe a method for obtaining torus phase parameters from high-dimensional neural activity data. For more details, see the following paper [43] [44]. The n-dimensional torus *T*^*n*^ is represented as *n* direct products of the unit circumference *S*^1^ as *T*^*n*^ = *S*^1^ ×*S*^1^ ×… *S*^1^. The next problem is to find the circular coordinates *θ* : *X*→ *S*^1^ that map from the VR complex *X* to *S*^1^. We use the parameter *f* to define the circular coordinates *θ* on the VR complex *X*. Let the cocycle be *α*∈ C^1^(X; ℝ), and the smooth circular coordinates *θ* are obtained by minimizing the following equation.

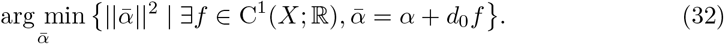

## Acknowledgments

The authors gratefully acknowledge Drs. Richard J. Gardner and Edvard I. Moser for permission to use the neurophysiological data [5].

