## Supplemental Fig for "Estimation of animal location from grid cell population activity using persistent cohomology": plos_kawahara_sup.pdf

### Supporting information

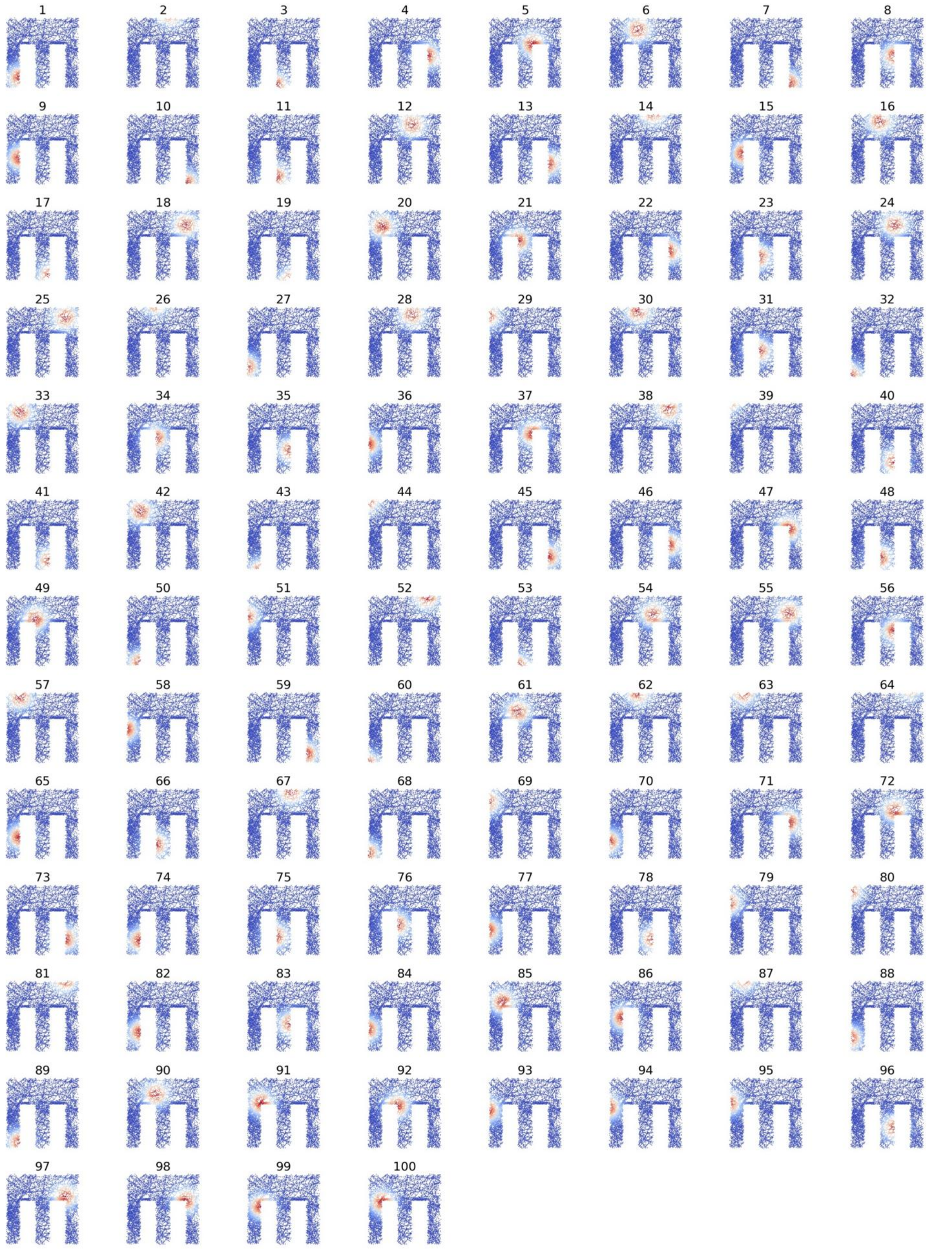


**S1 Fig. Receptive fields of 100 place cells.**

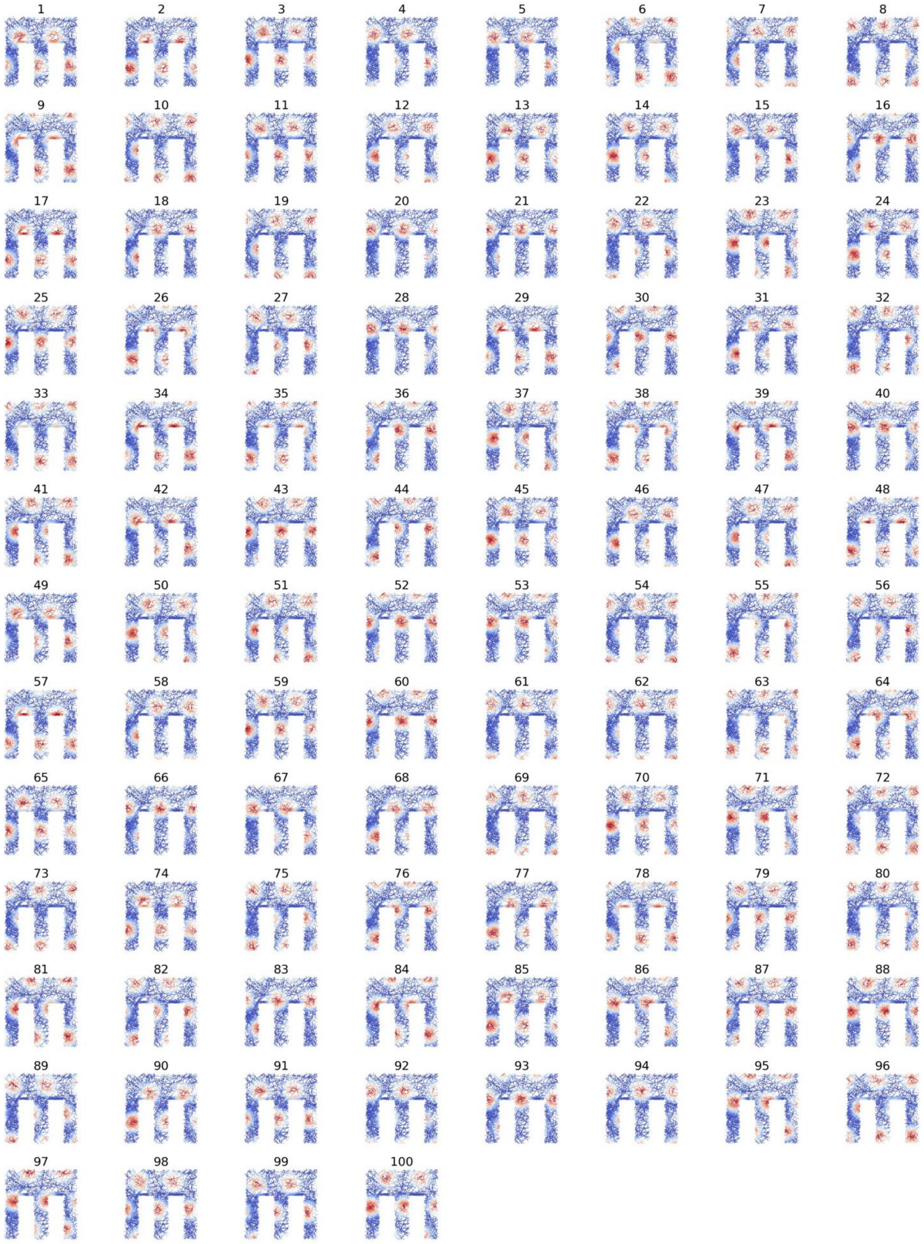

**S2 Fig. Receptive fields of 100 grid cells.**

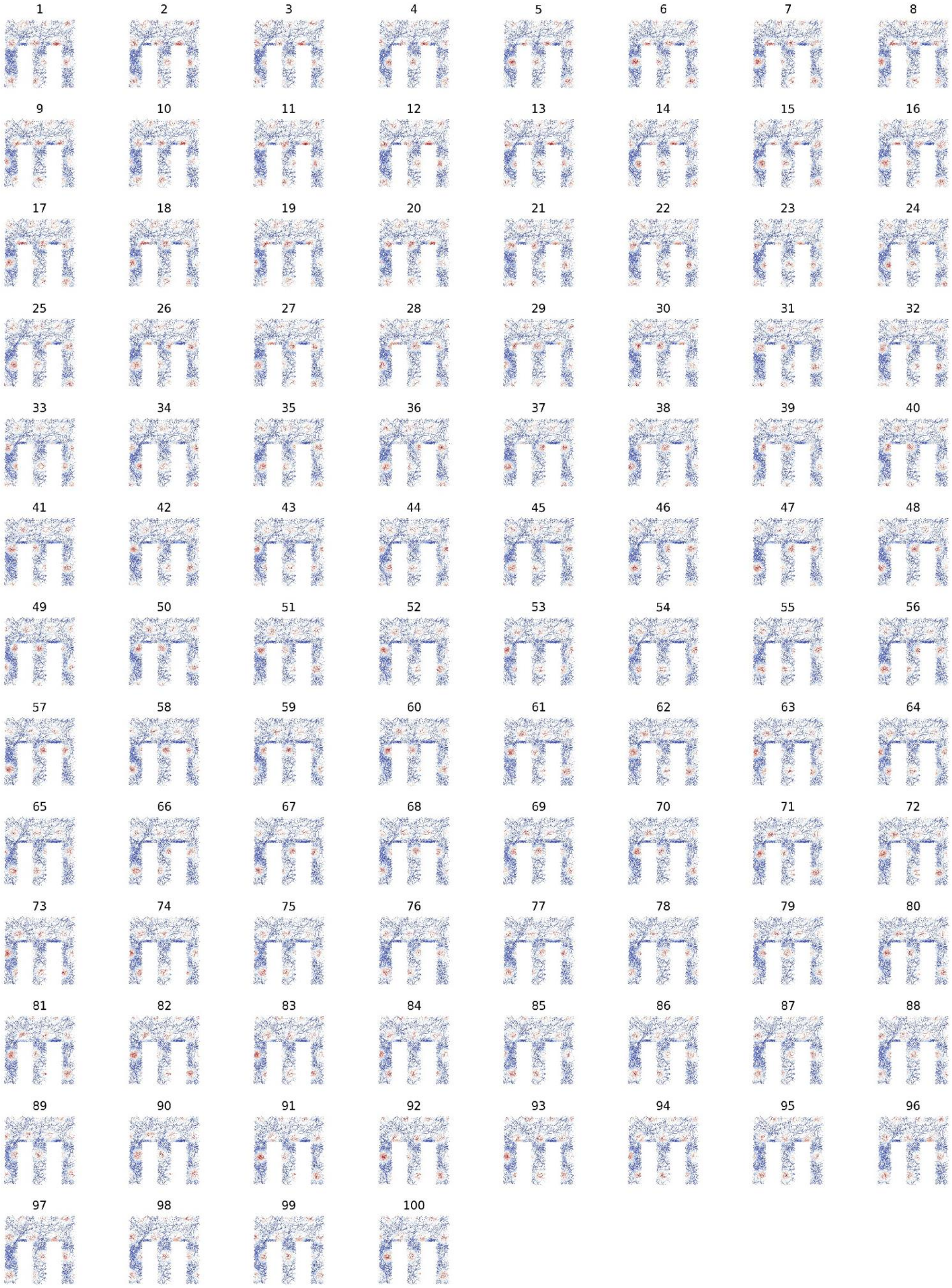


**S3 Fig. Receptive fields of 100 grid cells with small grid scale.**

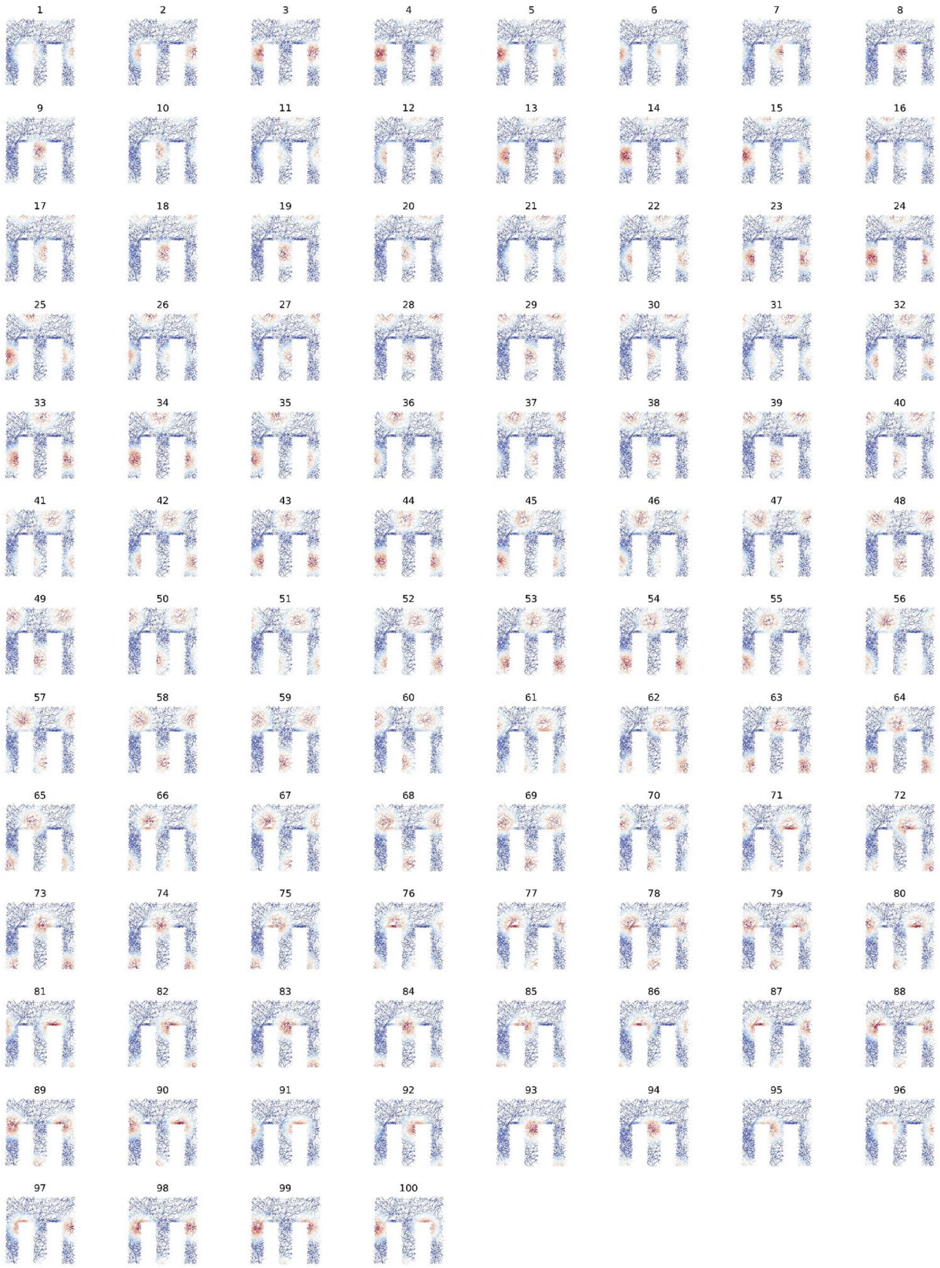


**S4 Fig. Receptive fields of 100 grid cells with large grid scale.**

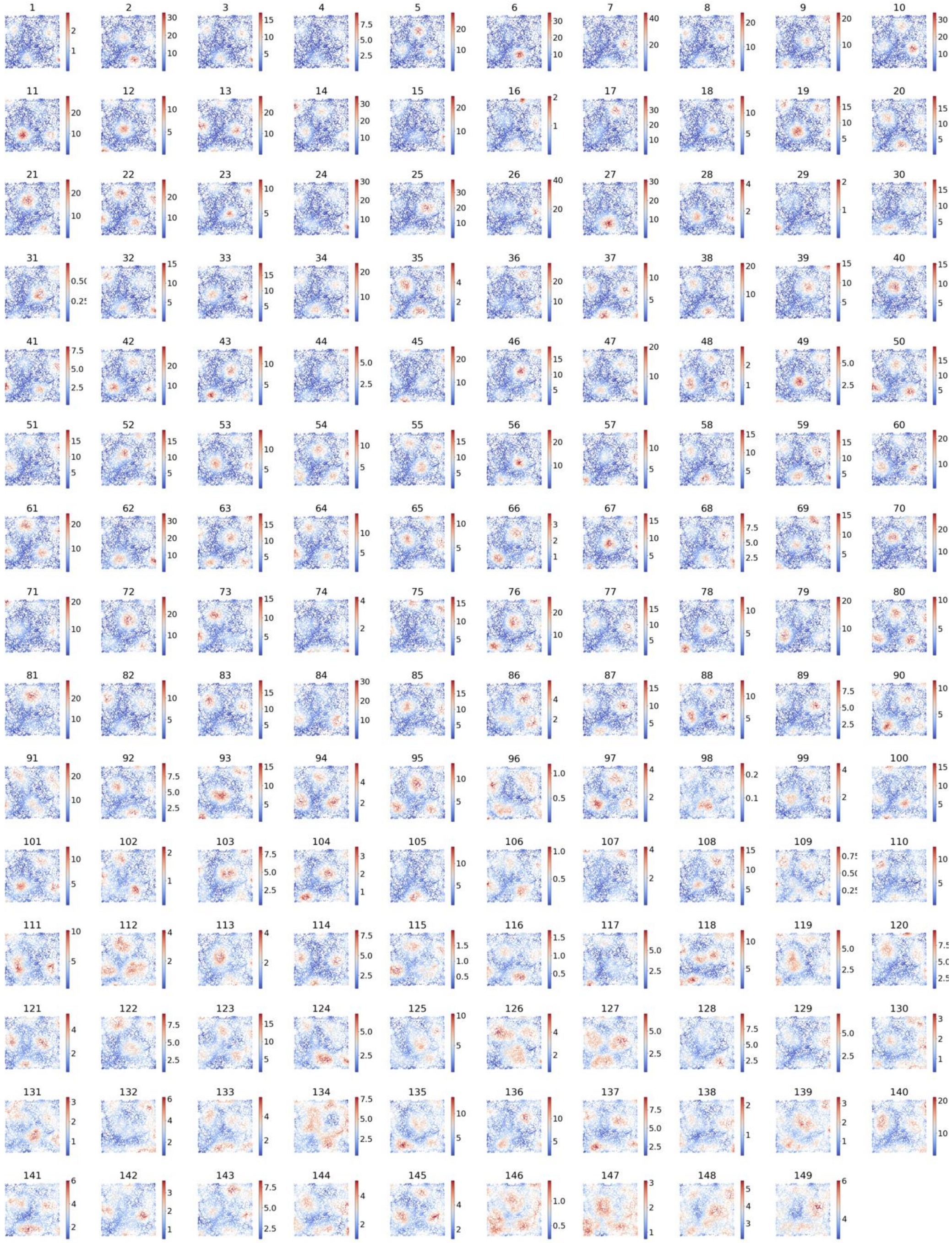


**S5 Fig. Receptive fields of 100 grid cells recorded from the rat's entorhinal cortex.**

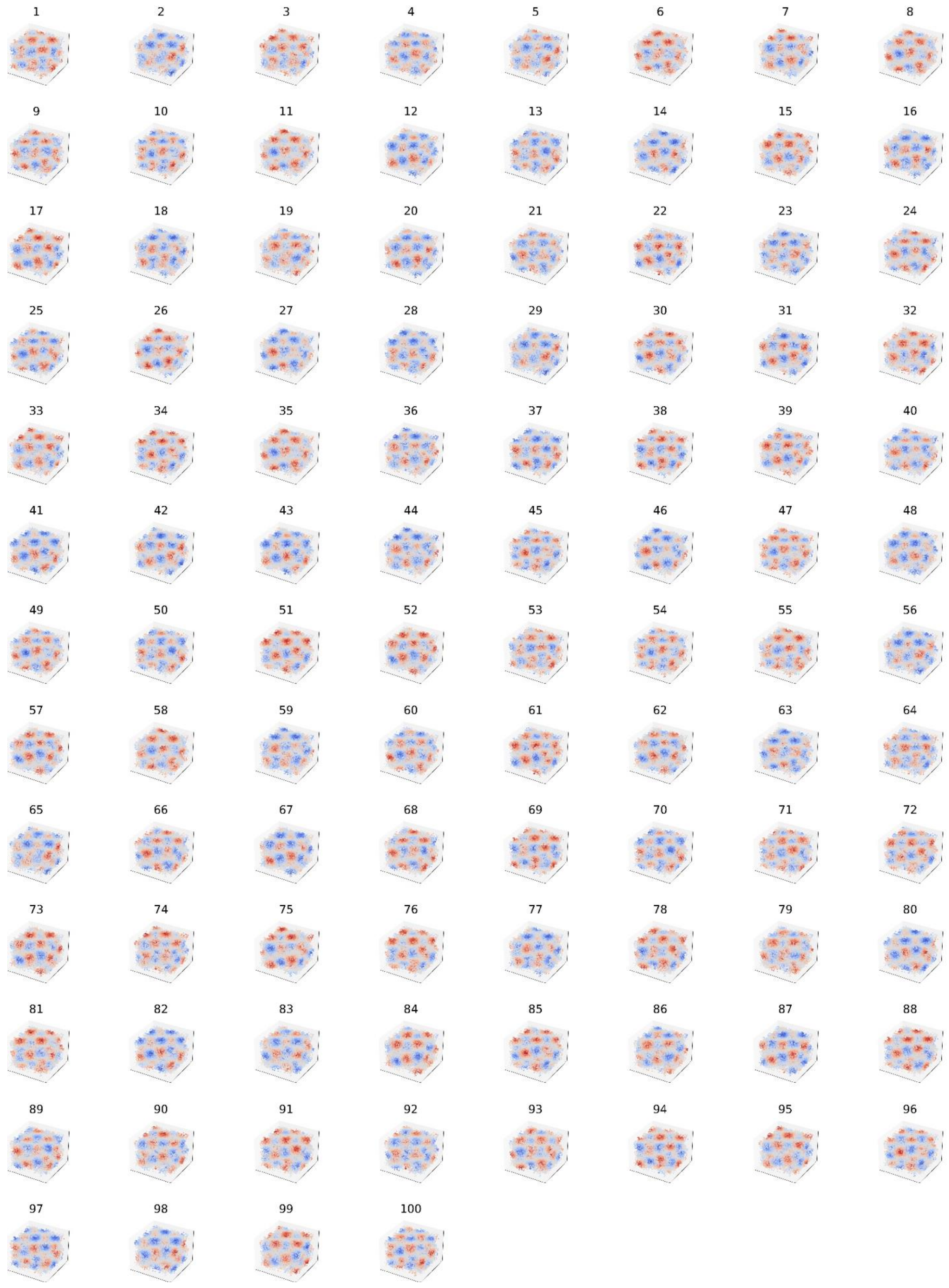


**S6 Fig. Receptive fields of grid cells in 3D-space.**

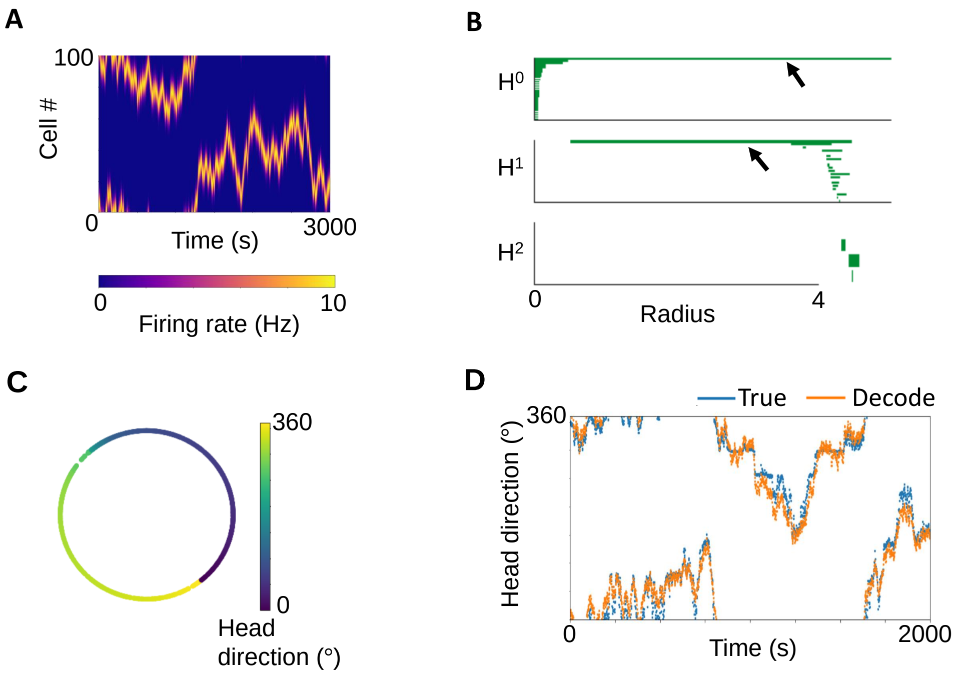


**S7 Fig. Estimation of the head direction in 1D-space using persistent cohomology.** (A) A raster plot of 100 head orientation cells for 3000 seconds. (B) Barcode. (C) Correspondence between the angle of head orientation at each time and the ring structure created by the neural activities. (D) Comparison of actual and estimated head direction.

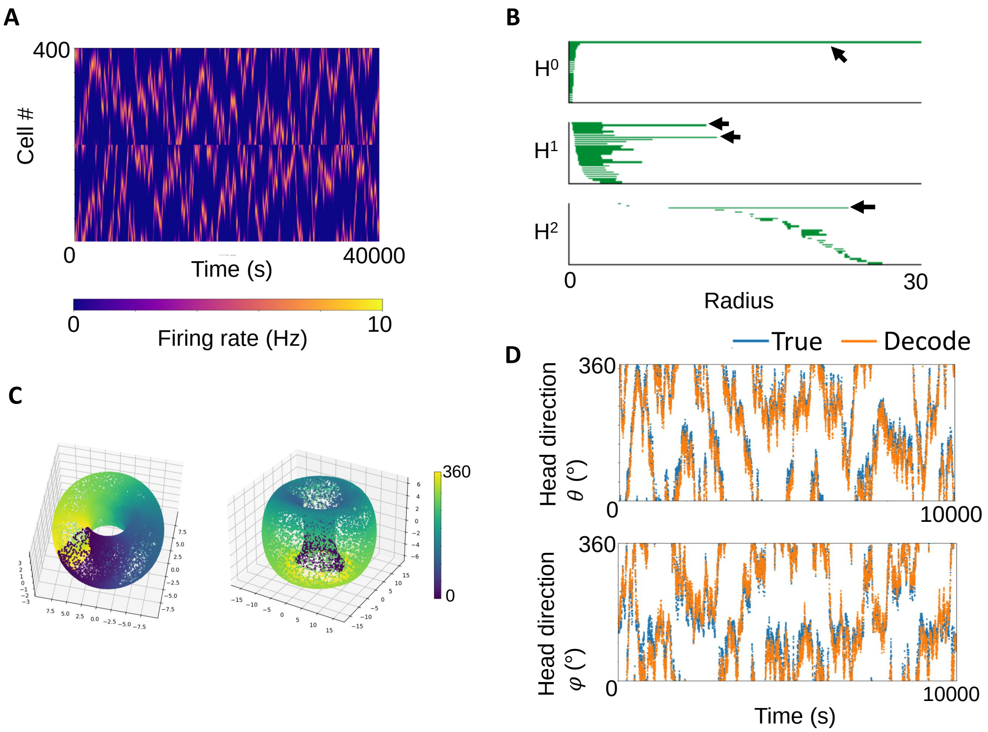


**S8 Fig. Estimation of the head direction in 2D-space using persistent cohomology.** (A) A raster plot of 400 head orientation cells for 4000 seconds. (B) Barcode (C) Correspondence between the angle of head orientation at each time and the 2-D torus structure created by the neural activities. (D) Comparison of actual and estimated head direction.
